# Peer presence increases adolescents’ prosocial behavior by speeding the evaluation of rewards for others

**DOI:** 10.1101/2021.03.17.435800

**Authors:** Nicolette J. Sullivan, Rosa Li, Scott A. Huettel

## Abstract

Peer presence can elicit maladaptive adolescent decision-making, potentially by increasing sensitivity to the rewards one receives. It remains unknown whether peer presence also increases adolescents’ sensitivity to others’ rewards, which could have an adaptive effect in contexts allowing pro-social behaviors. Here, we combine social utility modeling and real-time decision process modeling to characterize how peer presence alters adolescents’ processing of self and other rewards. We found that adolescents behaved selfishly when privately allocating rewards for themselves and a peer in an incentive-compatible task. In peer presence, however, adolescents became more altruistic. Real-time estimates of reward processing collected using computer mouse tracking showed that altruistic behavior was associated with relatively earlier processing of peer-rewards relative to self-rewards, such that peer presence sped peer-reward processing without altering self-reward processing. Our results indicate a causal mechanism through which peer presence prompts greater prosocial behavior by altering how adolescents process prosocial outcomes.

## Introduction

Public health data indicate that many maladaptive behaviors peak in adolescence (Preusser, Ferguson et al. 1998, Chen, Baker et al. 2000, Lundborg 2006, Simons-Morton and Farhat 2010, Zimring and Laqueur 2014), a phenomenon often attributed partially to an adolescent-specific heightened sensitivity to reward (Galvan 2010, Walker, Bell et al. 2017). Peer presence has been shown to exaggerate adolescents’ reward sensitivity and, consequently, their reckless behaviors (Gardner and Steinberg 2005, Chein, Albert et al. 2011, Smith, Chein et al. 2014, but see Braams, Davidow et al. 2019). Previous studies of peer influence on adolescent decision-making have thus far largely investigated the mechanisms underlying processing of outcomes specifically for self-rewards (Gardner and Steinberg 2005, Chein, Albert et al. 2011, Smith, Chein et al. 2014) or outcomes tied to self-rewards (Van Hoorn, Van Dijk et al. 2016, van Hoorn, van Dijk et al. 2016), yet many of adolescents’ everyday decisions affect not only themselves but also their peers (see Ciranka and van den Bos 2019 for a review). Therefore, it remains unknown whether and how the presence of a peer influences how adolescents process rewards for that observing peer *separately* from a more general effect on reward processing, and how this affects adolescents’ decisions when others’ outcomes are also at stake.

Here, we consider the hypothesis that the presence of a peer increases sensitivity to others’ rewards, which in turn can facilitate *prosocial* behavior. We evaluate a specific mechanism for this increased sensitivity: reductions in the relative time required for the brain to process other-related rewards. Our study is motivated by a robust set of decision process models, often collectively referred to as accumulator models, that have been shown to account for both choices and their response times across a broad range of choice domains (Busemeyer and Diederich 2002, Milosavljevic, Malmaud et al. 2010, Krajbich and Rangel 2011, Krajbich, Lu et al. 2012, Hutcherson, Bushong et al. 2015, Ratcliff, Smith et al. 2016, Fisher 2017). In these models, the relative evidence toward one response option over the other accumulates until it reaches a threshold and triggers a choice. Most such models share one assumption: option comparison cannot begin until the qualities (i.e., values or attributes) of each option have been estimated. This onset time has been found to vary according to several characteristics of choices and varies based on the salience of each option (Chen, Mihalas et al. 2013) and attribute types (Dshemuchadse, Scherbaum et al. 2013, Sullivan, Hutcherson et al. 2015). Accumulator models predict, and recent research has confirmed, that attributes processed earlier during option comparison receive a greater weight in choice, due to their relatively longer contribution to the decision process (Sullivan, Hutcherson et al. 2015, Lim, Penrod et al. 2018).

Under many conditions, potential reward outcomes for the self are likely to be more concrete and relevant to the decision maker than rewards a peer may receive (Liberman and Trope 2008). If so, self-rewards would be processed relatively earlier, resulting in decisions that favor advantageous inequity (i.e., where the self’s outcomes are better than the peer’s) and reject disadvantageous inequity (i.e. where the self’s outcomes are worse than the peer’s). We propose that peer presence alters this decision process by speeding the processing of peer rewards, either by increasing their salience or reducing their abstractness, shifting people toward more altruistic choices. This may be particularly true in adolescents, who are more sensitive to peers in general than adults (Gardner and Steinberg 2005, Logue, Chein et al. 2014) and spend more time with their peers as well (Csikszentmihalyi, Larson et al. 2014).

In the current study, adolescents made incentive-compatible decisions about self-other monetary divisions within a multi-round dictator game, both alone and while watched by a peer friend (Fig. 1; Forsythe, Horowitz et al. 1994). We quantified the influence of peer presence on social preferences using robust social utility models (Fehr and Schmidt 1999, Charness and Rabin 2002) that capture fine-grained differences in how self- and peer-rewards affect decision-making under conditions of advantageous and disadvantageous inequity. We tracked computer mouse movements during the choice process, which allowed us to pinpoint the times at which self and other rewards begin to contribute to decisions and how those times were altered depending on whether participants were alone or watched by a peer. The combination of social utility models and process tracing methods provides an integrated explanation for the paradoxical features of adolescent decision-making: both selfish decisions for oneself and altruistic decisions for one’s peers depend on the relative speed of self and other reward processing across decision contexts.

**Fig. 1.**
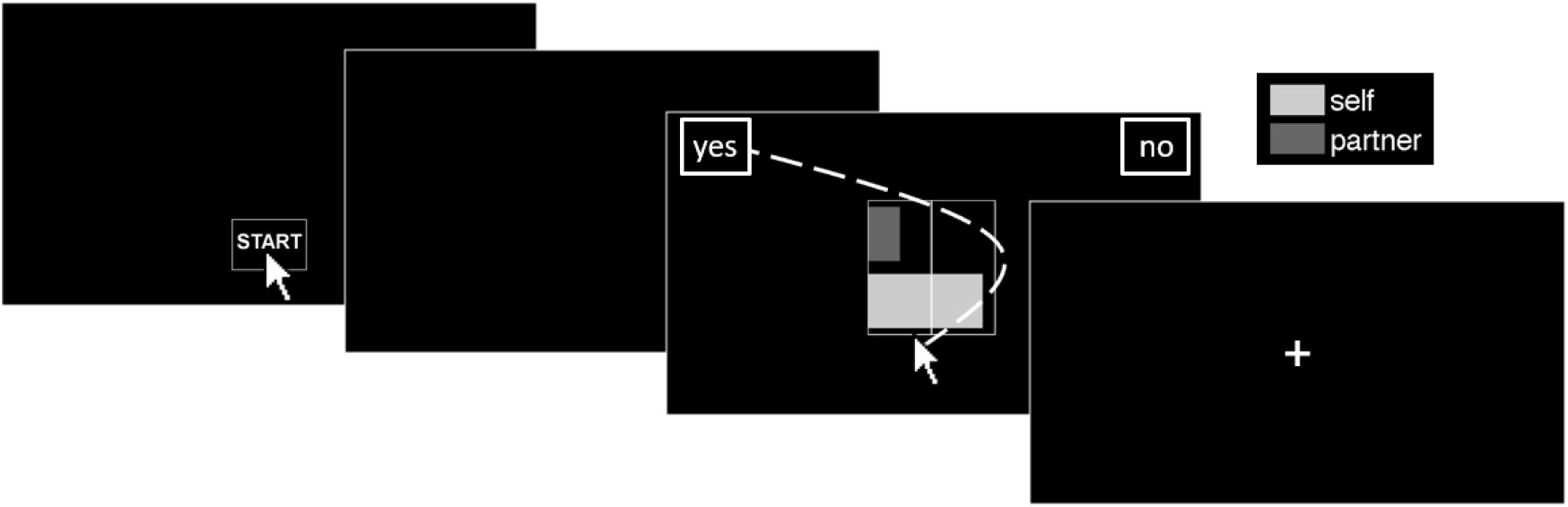
Reward allocation task. Participants used a computer mouse to accept or reject allocations of rewards for themselves (light grey bars) and their peer (dark grey bars; labeled as “partner” to the participant). Participants began each trial by clicking a START button. Once participants began moving the mouse, the current offer of between 0 and 10 points for self and 0 and 10 points for peer appeared on the center of the screen. Participants smoothly and continuously moved their mouse to the “YES” box to accept the displayed offer or the “NO” box to reject it and instead receive 5 points for self and 5 points for peer, represented by the vertical line drawn at the halfway mark of the graphical display. The trajectory of the mouse movement – and particularly, the inflection points of any change in that movement – provided information about the relative timing of when self and peer rewards influenced the decision process. Choices on randomly selected trials were converted to bonus payments for both the participant and the peer at the end of the session (see Methods).

## Methods

### Participants

Fifty-eight adolescents (U.S. high school juniors and seniors) were enrolled in the study; one participant who missed too many catch trials (see *Task* below) was excluded prior to data analysis. Thus, our final sample comprised 57 adolescents (mean age = 17.2 years; range = 15.7 to 18.8 years; SD = 0.56 years; 38 F) who completed the experimental conditions in a within-participant design, with a complementary adult sample of 29 participants reported on in the Supplement (see “Adult Alone comparison sample”). We used a cultural definition of adolescence such that all participants were still enrolled in high-school and had not yet reached full independence from their parents/guardians (Crone and Dahl 2012). Participants were drawn from a community participant pool. The adolescent sample is made up of self-reported ethnic identity of: 4 Hispanic or Latino, 53 non-Hispanic or Latino; 15 Asian, 7 Black or African American, 30 White or Caucasian, and 3 reporting more than one race. Two participants declined to respond to this question. The demographics of our sample generally matches that of North Carolina, apart from an overrepresentation of participants who self-identified as Asian.

All adolescents completed 300 trials in the Alone condition, and 55 of 57 adolescents also performed the task while being watched by their peer (mean: 288.3 trials, sd: 34.5 trials). Of the 55 adolescents who completed the Watched condition, 47 completed all 300 trials, while 8 completed as many trials as time allowed (range: 131-262 trials). Two dyads did not complete the third round of the task (i.e., the second *Watched* condition) because of time constraints.

The targeted sample size (60) was determined based on a pilot dataset that included a similar binary choice task. First, we calculated the difference in self and other latencies (the times at which self and other rewards are processed; see “Mouse tracking: Data Analysis” below) in a pilot dataset using the same task (N=32) which generated an approximate sample size of 50 (one-sample t-test of self vs. other latencies vs. 0, sampsizepwr function in MATLAB and a p < 0.05 threshold and power = 0.99). Because latencies are often non-normally distributed, we added 10 additional participants to this size estimate (Lehmann 1998).

All participants and parents/legal guardians of adolescent participants gave written informed consent in a protocol approved by the university’s Institutional Review Board. Note that the informed consent process occurred on a separate day, prior to the experimental sessions. Moreover, while participants were informed at the time of recruitment and consent of the possibility that their peer would observe them in the experiment, the task instructions provided no advance information about repeating the task in Watched and Alone conditions. Thus, participants were blinded to the experimental conditions until the moment of participation.

Adolescent participants enrolled in the study with a same-gendered peer who also completed the study; that is, there were 29 adolescent dyads who arrived to the testing site together and whose decisions influenced each other’s payments. Participants received instructions and completed 300 trials of the task in separate rooms (*Alone* condition). This was always performed first to assess baseline inequity aversion without the potential contaminating influence of the Watched condition. Next, both adolescents of the dyad were moved into the same testing room (*Watched* condition), and one adolescent completed another 300 trials while the peer watched from a nearby chair. Finally, if time permitted, adolescents in each dyad switched the watching/task roles for up to 300 more trials of the task. During the Watched condition, both participants were instructed not to communicate with each other, but the task-watcher was asked to record on a private worksheet how much he or she agreed with their peer’s choices. This procedure was implemented not only to ensure that the task-watcher remained engaged, but also to emphasize that the peer was watching (and evaluating) behavior in the task.

### Task

Participants used a computer mouse to accept or reject point allocations for themselves and their peer. Each trial began with the display of a box containing the word “START” at the bottom center of a black screen. When the participant clicked “START”, the screen went black. Once the participant began moving the mouse, the current offer appeared in the center and boxes containing “YES” and “NO” appeared in the top corners of the screen (randomly counterbalanced across trials). Participants were instructed to smoothly and continuously move their mouse to “YES” to accept the displayed offer or to “NO” to reject the displayed offer; if the offer was rejected, then they and their peer would each receive the default allocation of 5 points. Offers were depicted using colored bars of varying lengths to represent points for self (light grey) and points for peer (dark grey), with full bars representing 10 points (see Fig. 1). A vertical line was drawn at the halfway mark to represent the reference 5/5 allocation option. The top/bottom position of the self and peer bars was randomly counterbalanced across trials. Stimuli were presented using the Psychophysics Toolbox (Brainard 1997) for MATLAB.

Potential offers ranged from 0 to 10 points for self and 0 to 10 points for peer in half-point increments, save for the exclusion of 5/5 and 0/0 offers. For each participant, we drew a unique set of offers from this range such that a person of average preferences (as determined by pilot testing) would be close to indifferent on 1/3 of trials, readily accept 1/3 of trials, and readily reject 1/3 of trials. Trials in which self and peer had equal payouts were treated as catch trials, such that participants should always accept catch trials in which the self/peer payouts were greater than 5 (e.g., 8/8) and reject catch trials in which the self/peer payouts were less than 5 (e.g., 4/4). One adolescent’s data was excluded prior to analysis for not having enough correct catch trials to pass a binomial test of greater than chance performance. Catch trials were not included in subsequent analyses.

For incentive compatibility, one randomly selected trial from each completed run was be converted into dollars using a predetermined exchange rate that was revealed to the participant at the end of the session (38 cents/point if a participant completed two runs; 75 cents/point if a participant only completed one run). Thus, participants did not know the exact monetary value of the offers, which controlled for potential individual differences in valuation of money. Participants were paid based on the randomly selected trials that they completed and that their peer completed (e.g. paid as both “self” and as “peer”). This procedure was discussed with each participant before the experimental task.

### Choice models

We used maximum likelihood estimation to fit participants’ choices on each run to four different social utility models (see Supplementary Methods). For every completed trial, the value of that trial’s offer was set relative to the default offer (i.e., 5-self, 5-peer) by subtracting 5 from the values of what was offered for self and for peer. The value of the rejection was thus set to 0. A sigmoid function (Eq. 1) translated the value difference between acceptance and rejection into a probability of acceptance. The temperature parameter μ represents the sensitivity (steepness) of the sigmoid function, or with higher values indicating greater choice consistency.

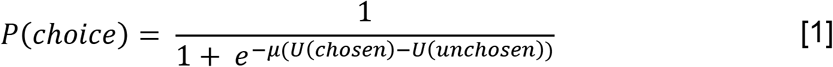

We found that the model from (Charness and Rabin 2002) yielded the lowest AIC and BIC values and was therefore used in all subsequent analyses (Eq. 2):

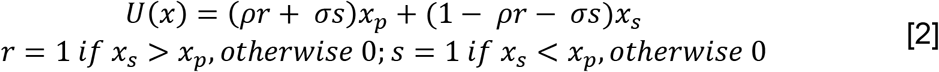

Within this model, the parameter ρ describes preferences when reward for self is greater than reward for peer, while σ describes preferences when reward for self is less than reward for peer. For both ρ and σ, larger values indicate an increasing relative decision weight on rewards for the peer – with less than 0.5 indicating greater weighting on self-rewards, 0.5 indicating equal weighting for self- and peer- rewards, and greater than 0.5 indicating greater weighting on peer-rewards. Our model-fitting procedure searched the parameter space between −10 and 10; most participants’ parameter estimates fit between 0 and 1. Negative values indicate that participants lose utility for each additional unit of peer-reward, while values greater than 1 indicate that participants lose utility for each additional unit of self-reward.

### Mouse tracking: Data Collection and Processing

The cursor’s position was tracked with a temporal resolution equal to the screen refresh rate (60 Hz). At each time point, the cursor angle was normalized such that 0° represented direct upward movements, −45° represented movements directly toward the left option, and 45° represented movements directly toward the right option.

To ensure that we analyzed data from trials on which subjects made continuous, goal-directed mouse movements, we established two *a priori* exclusion criteria: trials with reaction times (RTs) greater than two standard deviations above a subject’s mean, and trials in which the mouse trajectory crossed the midline of the x-axis (i.e. an invisible vertical line bisecting the screen) more than three times (a mean of 9.2% excluded trials per subject).

### Mouse tracking: Data Analysis

We used multiple linear regression to estimate (for each participant and condition independently) how self (*S*) and peer (*P*) rewards influenced the cursor angle at each time point during the mouse movement. For timepoint t, this regression (Eq. 3) used the differences in rewards associated with the left and right options to predict the mouse cursor’s angle of movement between time t and t+1, Θ*_t_*.

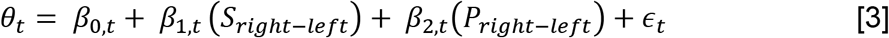

Next, we normalized each timepoint’s self and peer coefficients from Eq. 3 to be proportional to the coefficient’s full final weight, β_N_ (Eq. 4).

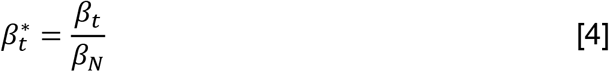

Finally, we identified the timepoint at which this β*t for each attribute became greater than zero; that is, when that attribute began to gain some portion of its final attribute weighting. To do this, we modeled the increase in proportion of final attribute weight on cursor using a generalized logistic function, using a population growth equation (e.g., Richards 1959, Zwietering, Jongenburger et al. 1990, Tsoularis and Wallace 2002). Unlike a traditional growth model, our function begins at zero, so the model was modified to a piecewise function (Eq. 5). This function includes a parameter (t*) that specifies the time begins (i.e., the time at which the function changes from β*_t_ = 0 to β*_t_ > 0). We use this as our estimate of the time at which an attribute begins to influence the mouse cursor, and therefore the time at least by which the brain must have processed this value information, which we label as *attribute latency*. This extensive procedure was implemented to minimize the possibility that the final timepoint’s reward coefficient – that is, the weight participants placed on self- or peer-rewards – did not artificially bias latency estimation (see reference Sullivan, Hutcherson et al. 2015 for further discussion of this concern). However, we should also note that all results hold using the uncorrected estimation time used in (Sullivan, Hutcherson et al. 2015). Processing speeds for self and peer rewards were unable to be estimated for 11 and 12 participants respectively, so those participants were excluded from analyses of processing speeds.

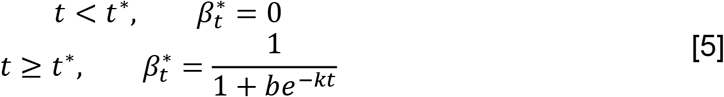

## Results

We tested our hypotheses using an incentive-compatible, repeated-trial dictator game in which participants chose between two allocations of rewards for themselves and a peer friend: an on-screen allocation that varied across trials (i.e., could be larger or smaller, or better for the participant or the friend) or a reference allocation in which they and their peer received identical, intermediate rewards (Fig. 1). Choices were made either alone, or while their peer watched their choices. To indicate whether they preferred the on-screen division or the reference option, participants moved their computer mouse cursor from its starting position at the bottom of the screen to one of the response boxes on the top left and right of the screen. Some trials presented a situation of “advantageous inequity,” in which the decider was to receive more points than their peer. Other trials presented a “disadvantageous inequity” in which the decided would receive less than their peer. Participants were not told the conversion rate between points and monetary amounts until after the task.

### Peer presence shifts tolerance for inequity

Adolescents generally accepted offers involving increased rewards to both themselves and their peer, relative to rejecting the offer in favor of 5-self/5-peer in both the Alone and Watched conditions (Figure 2A, B; top right quadrant). They also generally rejected offers in which both self and peer would receive fewer than 5 points in both conditions (bottom left quadrant). The presence of a peer also led to decreased acceptance of allocations involving *advantageous* inequity (i.e., receiving more than their peer) but increased acceptance of allocations involving *disadvantageous* inequity (i.e., receiving less than their peer; Figure 2C). This difference by condition was particularly striking for decisions involving advantageous inequity when the participant would receive more, and their peer less, than the equitable off-screen reference of 5-self/5-peer (top left quadrant).

To formally characterize participants’ social preferences, we fit four social utility models (see Methods & SI). The best-fitting model (by Charness and Rabin; see Charness and Rabin 2002) captures two aspects of choice behavior: relative weighting for self and peer reward when self earns more than peer (advantageous inequity; parameter ρ), and relative weighting for self and peer reward when self earns less than peer (disadvantageous inequity; parameter σ). In either case, the key parameter ρ or σ fits the relative weight placed on self (values < 0.5) versus peer rewards (values > 0.5), with parameters of 0.5 indicating equal weight placed on self and peer rewards. We allowed our model to fit below zero and above 1, meaning that participants could lose utility per unit of either peer (weights below 0) or self (weights above 1) rewards.

When alone, adolescents placed a larger weight on self- than peer-rewards under advantageous inequity (mean ρ = 0.35) and placed all weight on self-rewards with no weight on peer-rewards under disadvantageous inequity (mean σ = −0.05; Fig. 2D; ρ vs. σ paired Wilcoxon signed rank test, d = 0.58, W = 1249, z = 3.36, *p* < 0.001). Five participants’ behavior was poorly captured by this model, resulting in outlier parameter fits that fell outside the display range so are excluded from Fig. 2D (see Fig. S1 for a figure with all participants). Bar means plotted on Fig. 2D and all statistics, however, include these participants.

**Fig. 2.**
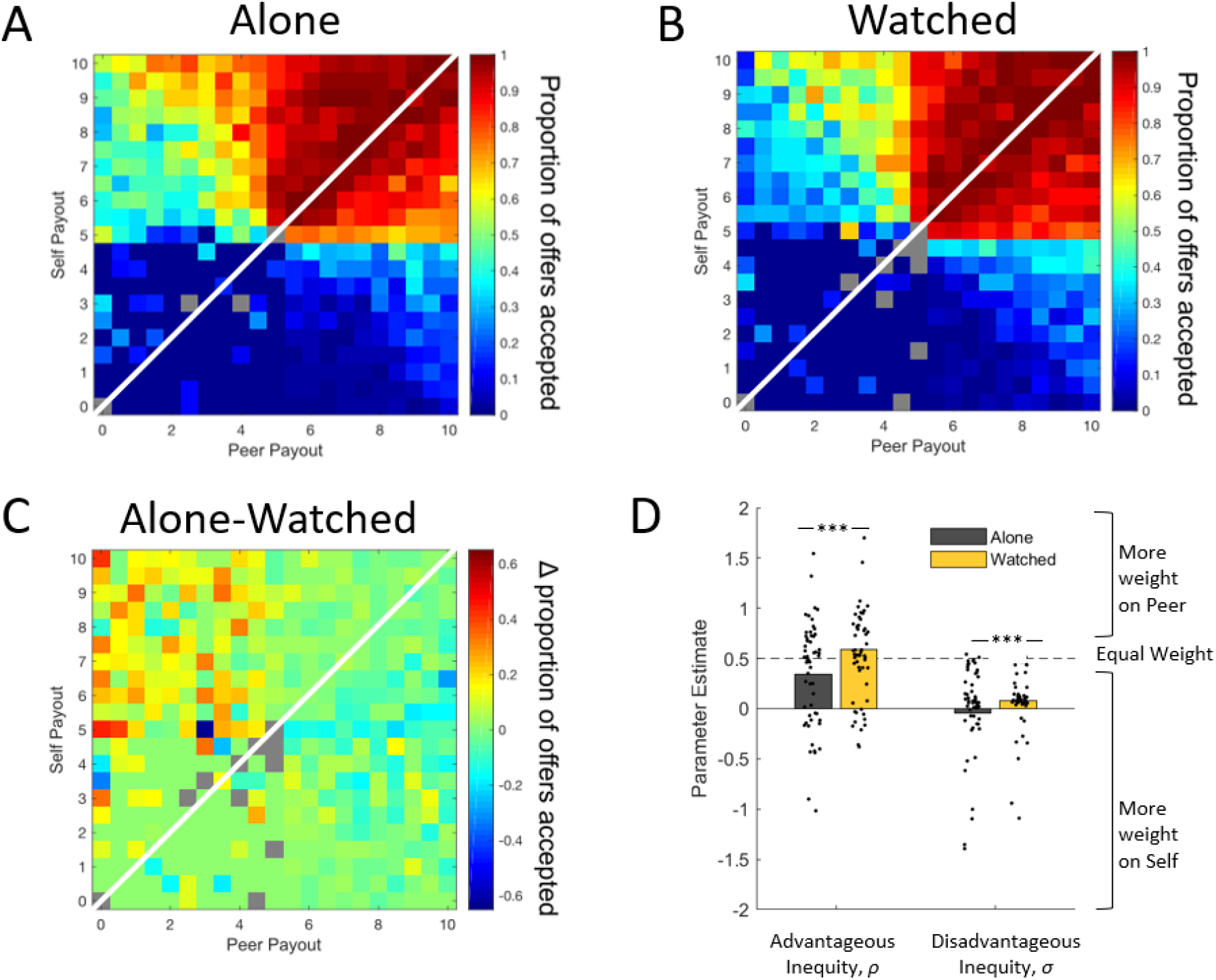
Adolescents more averse to advantageous inequity when watched. Shown are the proportions of accepted offers for each combination of self and peer payouts when adolescents completed the task (A) alone and (B) while watched by a peer. Rejecting the offer resulted in a payout of 5-self/5-peer. Panel (C) depicts the change in offer acceptance when moving from completing the task Alone to being watched by a peer. Warm colors represent more offers accepted in the Alone condition, and cooler colors represent more offers accepted in the Watched condition. For all three heatmaps, grey tiles represent offers shown fewer than three times across all participants. The white diagonal line represents a division between situations of advantageous inequity (upper left triangle) and disadvantageous inequity (lower right triangle). (D) Average inequity parameters when alone and when watched by a peer. Adolescents exhibit significantly greater ρs and σs when watched compared to when alone, indicating greater weight on peer rewards (Xp) and less weight on self rewards (Xs). Dots represent individual participants. For display purposes, the individual dots for the six participants whose parameters fit outside of the [-2, 2] range are included in the mean bars, but not displayed. *** indicates p < 0.001.

Next, we evaluated how peer presence altered prosociality. When watched, adolescents showed increased aversion both to advantageous inequity (mean ρ = 0.59) and to disadvantageous inequity (mean σ = 0.08), indicating that adolescents place greater weight on peer-reward when a peer is present (Fig. 2). Because the distributions of ρ and σ estimates were non-normal (ρ: skewness = 0.86, kurtosis = 13.42, Shapiro-Wilks W = 0.80, *p* < 0.001; σ: skewness = −2.93, kurtosis = 14.65, Shapiro-Wilks W = 0.70, *p* < 0.001), we used the non-parametric paired Wilcoxon signed rank test to determine that the increase in ρ and σ parameters when Watched was statistically significant within the subset of 55 participants who completed both the Alone and Watched conditions (ρ: d = −0.37, W = 337, z = −3.63, *p* < 0.001; σ: d = −0.24, W = 289, z = −4.03, *p* < 0.001).

To assess baseline inequity aversion without possible contaminating or priming effects of being watched by a peer, the Alone condition was always performed first. This does not allow us to control for task order and practice effects. However, due to the nature of the Watched condition, some participants performed the Watched task first while their partner watched, and others performed it second after being watched by their partner (randomly determined). By assessing changes in inequity aversion by whether the participant was the decider first or last in the Watched task, we can gain some insight into whether practice effects influence inequity aversion. Instead, we found that there was no statistically significant difference in either ρ or σ as a function of completing the Watched task first or last (ρ medians first = 0.65, last = 0.52; d = 0.19, U = 700.00, z = −0.51, p = 0.61; σ medians first = 0.04, last = 0.10; d = −0.13, U = 760.00, z = 0.54, p = 0.59). This indicates that, at least in the Watched condition, practice effects are unlikely to be a significant driver of the inequity aversion observed in this study.

### Rewards to self are processed earlier than peer rewards when alone

We next used the mouse cursor position data to estimate the time point in the decision process where self-rewards and peer-rewards began influencing decision processes; these time points are known as *attribute latencies* (see Sullivan, Hutcherson et al. 2015 for method). To minimize any bias in this estimation that may arise from different amplitudes of the decision weights for these rewards, we normalized all time points to a proportion of each attribute’s full final amplitude, which results in a time series of each attribute’s influence on mouse movement that controls for each attribute’s final decision weight (Fig. S2; see Methods and SI for details).

We then estimated a piecewise growth model for each attribute that fits a parameter for attribute latency (i.e., the time point at which each curve in Fig. S2 diverges from zero). Non-parametric tests were used to compare latencies because self and peer latency distributions were non-normal (self latencies, skewness = −0.20, kurtosis = 3.96, Shapiro-Wilks W = 0.96, *p* = 0.01; peer latencies, skewness = 1.26, kurtosis = 5.68, Shapiro-Wilks W = 0.92, *p* < 0.001). In the Alone condition, information about self-rewards was processed earlier than information about peer-rewards (medians, self = 672 ms, peer = 933 ms, d = −0.82, W = 119, z = −3.36, *p* < 0.001). This suggests that, on average, rewards to the self are processed approximately 273 ms earlier than rewards to peers when adolescents are alone (the condition that is the standard set-up for most dictator-style monetary allocation games in the lab). Comparison to a control group of adults (see Supplementary Information) revealed that adolescents’ self-reward latencies are statistically indistinguishable from adults’ self- and peer-reward latencies. This indicates that adolescents’ slower processing of peer-rewards is specific to adolescence and not an artifact of our experimental procedures.

### Peer presence increases processing speed for peers’ rewards

Compared to the Alone condition, peer presence in the Watched condition significantly reduced the latency of processing information about rewards for peers by approximately 240 ms (Fig. 3, S3; medians, peer alone = 933, peer watched = 726; d = 0.57, W = 618, z = 3.18, *p* = 0.001). Peer presence, in fact, reduced the temporal advantage held by self-rewards in the alone condition to a non-significant difference. That is, self and peer latencies went from being statistically significantly different in the Alone condition (*p* < 0.001, see previous section) to not statistically significantly different in the Watched condition (medians, self watched = 615 ms, peer watched = 726 ms; d = −0.43, W = 248, z = −1.77, *p* = 0.08). Peer presence had no significant effect upon self-reward latency (medians, alone = 668 ms, watched = 615 ms; d = 0.27, W = 328, z = 0.85, *p* = 0.40). There was not a significant attribute-by-condition interaction (ANOVA interaction F(1,177) = 3.14, *p* = 0.08).

**Fig. 3.**
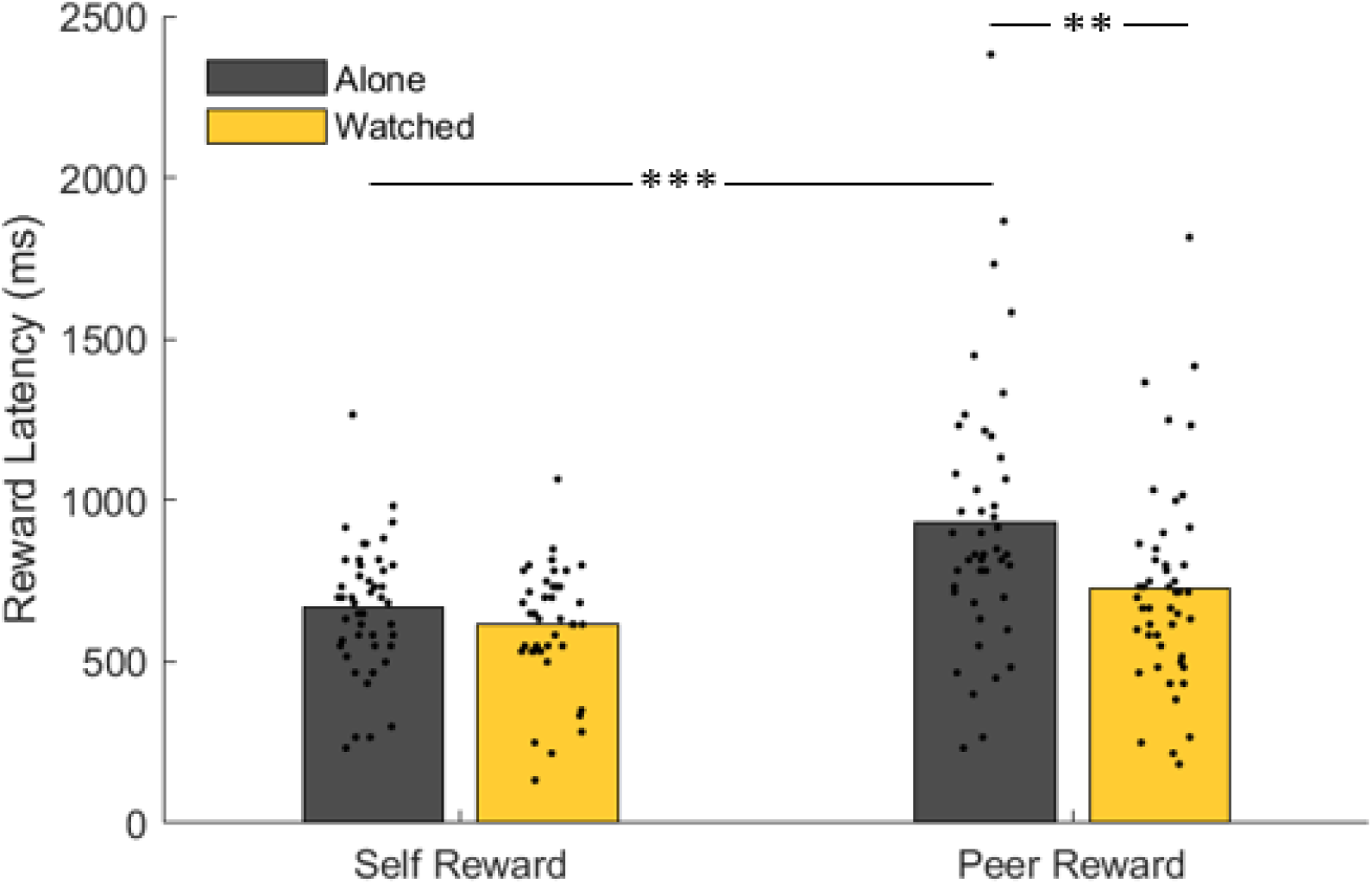
Results of reward latency estimation. Comparison of self and peer reward latencies in the alone and watched conditions. Dots represent individual participants. ** indicates p < 0.005, *** indicates p < 0.001.

### Faster relative peer processing was associated with prosocial behavior

We hypothesized that differences in attribute processing speed would ramify into participants’ expressed choices, because earlier-processed attributes can influence the decision process for a longer time. We refer to this earlier processing latency as a computational advantage (Sullivan, Hutcherson et al. 2015) – and we examined its relationship to the model parameters (Charness and Rabin 2002) for advantageous (ρ) and disadvantageous inequity (σ). Across our adolescent participants, we observed a strong relationship between the computational advantage of self-rewards and prosociality. Specifically, we found that the computational advantage for self-rewards was negatively correlated with prosociality under advantageous inequity (Fig. 4; linear regression r = 0.31, slope = 0.0005, 95% CI = [2×10^−4^ 0.001], *p* = 0.007). This relationship holds when excluding the outlier visible in Fig. 4 with a ρ value greater than 3 (*p* = 0.002). The computational advantage for self-rewards is not related to prosociality under disadvantageous inequity (Fig. S5; linear regression r = 0.05, slope = 5×10^−5^, 95% CI = [−2×10^−4^ 3×10^−4^], *p* = 0.67). These results provide a mechanistic explanation for prosocial allocation behavior: relatively faster processing of peer rewards leads both to greater decision weight for peer outcomes and increased aversion to advantageous inequity.

**Fig. 4.**
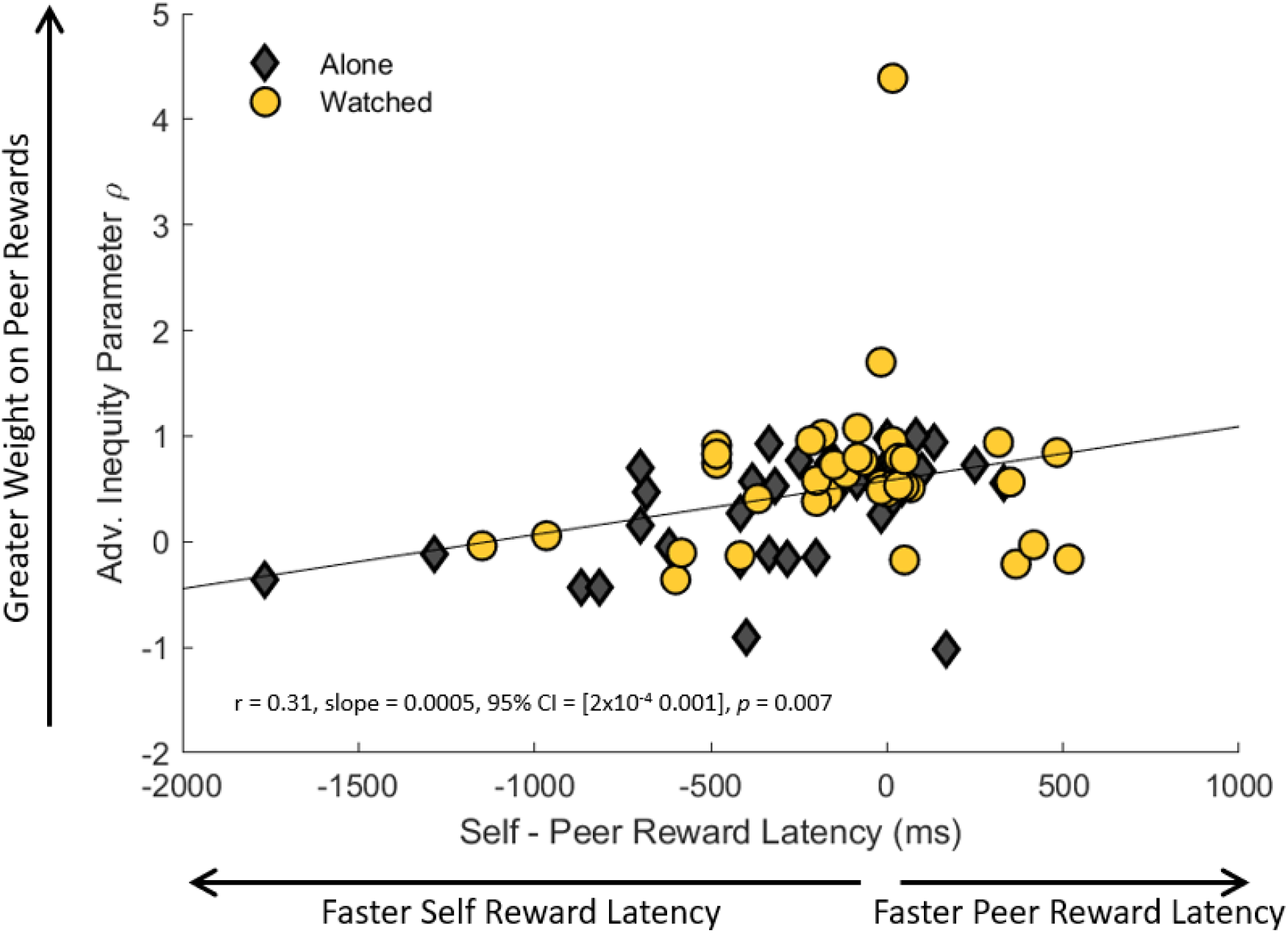
Relationship between advantageous inequity parameter ρ and the computational advantage of self-reward information. The markers depict self, relative to peer, processing speeds as a function of individual best-fitting advantageous inequity parameter ρ, with grey diamonds representing the alone condition and yellow circles representing the watched condition. The line represents the best-fitting linear regression line.

## Discussion

This study combines incentive-compatible economic games, process tracing methods, and computational modeling to characterize how peer presence influences prosocial behavior in adolescents. We found that that peer presence did not have a generalized effect on reward processing. Instead, its effects were selective: a greater influence on prosocial behavior under advantageous inequity than disadvantageous inequity and speeding only of rewards for peers but not for the self. These effects on processing measures had consequences for choices, such that the earlier that peers’ rewards began to influence the decision process relative to self-rewards, the more likely an adolescent was to consider peer payoffs in their choices. Together, modeling social utility through choice and processing speed through mouse tracking allowed us not only to dissociate the specific conditions in which peer presence alters adolescent behavior, but also to identify a potential mechanism for this behavior change. Compared to aggregate choice metrics, the use of social utility models provided a more fine-grained measure of other-regarding preferences in adolescents (Sul, Güroğlu et al. 2017). To our knowledge, this study represents the first demonstration of how *social* utility preferences of adolescents change in the presence of a peer. It is also the first application of mouse tracking to show that the speed of reward processing shapes pro-social tendencies.

Significantly, we found that peer presence had no effect on the time at which self-rewards entered the option comparison process – a surprising result considering that previous research on peer presence in adolescence would suggest an increased sensitivity to rewards more globally. There are several possible explanations for this finding. It is possible that the absence of significant change between conditions reflects a “floor effect,” such that self rewards are processed as fast as possible regardless of choice context. Alternatively, the effect of peers on adolescent reward sensitivity could be context-specific, meaning that peer presence alone was insufficient to heighten sensitivity to self-rewards. This latter explanation would reflect previous findings that adolescents are particularly sensitive to social contexts (Blakemore and Mills 2014). Future studies should investigate whether relative processing speed (and subsequent choices) can also be shifted in other ways, such as by increasing the salience of self- or peer-rewards through attentional or priming manipulations.

In this study, adolescents earned self- and peer-rewards while being watched by the same peer who would be affected by their decisions, as is often the case in real-world peer interactions. However, this means that our results cannot differentiate between two possible alternatives: one, that peer presence increases prosociality and speeds processing of peer rewards generally, or two, that peer presence only influences processing of rewards for the peer that is present (Diane and Richa 2013). However, a recent study found that young adults are more generous to both strangers and close friends when being watched by non-recipient others (van de Groep, Zanolie et al. 2020), providing evidence for the former interpretation. Our study design also cannot determine whether faster processing of peer rewards is driven by greater concern for the peer’s rewards themselves, or by anticipated social rewards for the self that may be received from seeming to behave more altruistically. Future studies could disentangle these psychological motivations for giving to others by allowing participants to remain anonymous or receive credit while making observed social decisions.

Early studies of adolescent susceptibility to peer influence generally focused on how peer presence prompted maladaptive behaviors like risk-taking (Gardner and Steinberg 2005, Chein, Albert et al. 2011, Haddad, Harrison et al. 2014, Smith, Chein et al. 2014). Here, we find that peer presence instead can have a salutary effect on adolescents’ behavior. This happens specifically in the context of altruistic choice, in which peers are more averse to inequity in the presence of an affected peer. With access to opportunities to engage in prosocial behavior, the presence of peers can be leveraged as a force for societal good, such as participating in group volunteering projects or engaging in political advocacy with peers (Middaugh, Clark et al. 2017).

## Supplementary Materials

### Supplementary Methods

#### Adult Alone comparison sample

Twenty-nine adult participants drawn from the Duke University community enrolled in the study with a same-gendered peer who completed a separate study. One adult was excluded from the study for missed catch trials, resulting in a final sample of 28 adults (mean age 22.7 years; range = 18.3 to 28.3; SD = 0.56; 12F; 1 Hispanic or Latino, 27 non-Hispanic or Latino; 12 Asian, 2 Black or African American, 13 White or Caucasian, 1 more than one race) who each completed a single 300-trial run Alone, as well as a demographics questionnaire. We note that though there was some age overlap in our two participant groups, we chose to use a cultural definition of adolescence by labeling 18-year-old high-school students as adolescents and those 18-years-old and older who live independently from their parents as adults (Crone and Dahl 2012).

### Supplementary Results

#### Model Comparison

We fit choice data to 4 different social utility models. Each social utility model is briefly described below, where xs denotes the payout for self, xp denotes the payout for peer, and U(x) denotes the participant’s utility for the trial.

##### Charness-Rabin Model

As discussed in (Charness and Rabin 2002), an individual’s utility is a function of two parameters, ρ and σ, that represent the relative valuation of self- and peer-rewards under conditions of advantageous inequity (ρ) and disadvantageous inequity (σ). When ρ or σ = 0, an individual is purely self-interested (i.e. receives no utility from the peer’s reward), when ρ or σ = 0.5, self and peer payouts are equally valued, when ρ or σ < 0.5, self-rewards bring more utility than equivalent peer-rewards, and when ρ or σ > 0.5, peer-rewards bring more utility than equivalent self-rewards.:

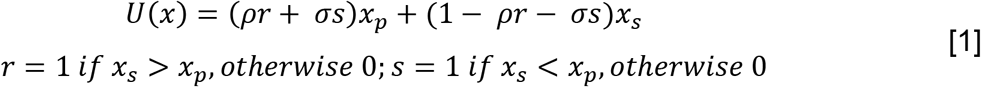

##### Fehr-Schmidt Model

As discussed in (Fehr and Schmidt 1999), an individual’s utility is a function of his/her own payout, advantageous inequity aversion (β), and disadvantageous inequity aversion (α). This model assumes that people dislike disadvantageous inequity more than they dislike advantageous inequity, and that people dislike advantageous inequity to a limited extent:

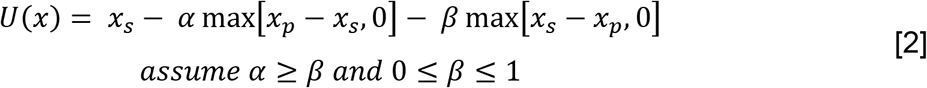

##### Unbounded Fehr-Schmidt Model

The Fehr-Schmidt model’s assumption that disadvantageous inequity aversion is always higher than advantageous inequity aversion was initially developed for cases in which participants made choices for unknown strangers. Because our participants played for known peers, this model removes the restriction that people dislike disadvantageous inequity more than they dislike advantageous inequity and only restricted both types of inequity aversion to be positive:

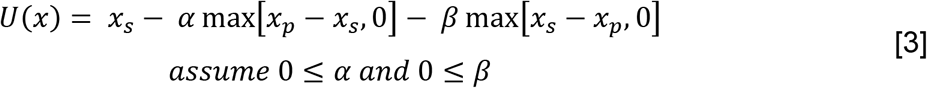

##### Unbounded Fehr-Schmidt Model with peer parameter

This model adds a parameter (γ) to the Unbounded Fehr-Schmidt Model to account for the peer’s raw payout:

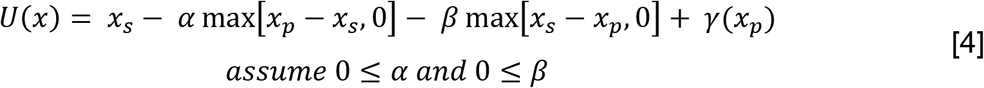

#### Adolescent Self- and Peer-Reward Latency Compared to Young Adult Control Group

For the young adult control group alone, self- and peer-reward latencies did not differ (median self-reward 741 ms, peer-reward, 687 ms; d = 0. 32, U = 702.50, z = 0.45, p = 0.66). Adolescents and the control group of young adults exhibited statistically indistinguishable processing speed for rewards for self (adolescent median = 668 ms, young adult median = 670 ms; d = −0.32, z = −0.48, p = 0.63). However, adolescents were markedly slower than adults at processing peers’ rewards (adolescent median = 933 ms, young adult median = 687 ms; d = 0. 61, U = 478.50, z = 3.09, p = 0.002).

#### Faster relative processing of peer-rewards associated with larger decision weights for peer rewards

For each adolescent participant, we estimated the weight participants placed on self and peer rewards, without imposing any additional model assumptions. To do so, we estimated decision weights for the influence of self and peer rewards (right–left) on choice (1= right, 0 = left), with one logistic regression per participant. Next, we estimated the relationship between the relative weight of self and peer rewards (self-peer) on relative self and peer reward latencies (self–peer). We found that earlier processing of peer, relative to self, latencies were associated with a larger weight placed on peer, relative to self, rewards in choice (Fig. S4. linear regression r = 0.44, slope = −0.0009, 95% CI = [−2×10^−3^ −3×10^−4^] *p* = 0.005).

---

**Fig. S1.**
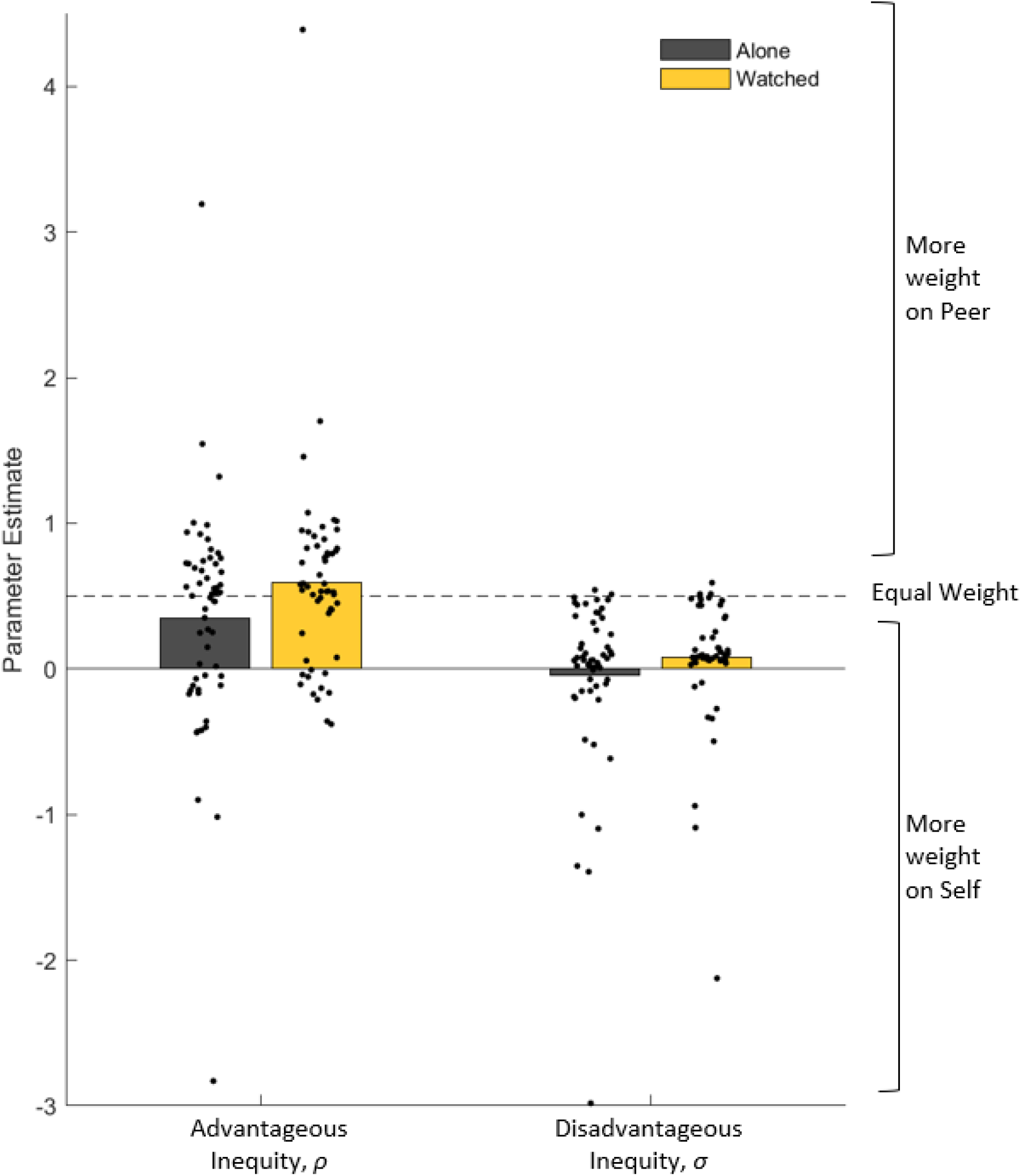
Average inequity parameters when alone and when watched by a peer. Adolescents exhibit significantly greater ρs and σs when watched compared to when alone, indicating greater weight on peer rewards and less weight on self rewards. Dots represent individual participants.

**Fig. S2.**
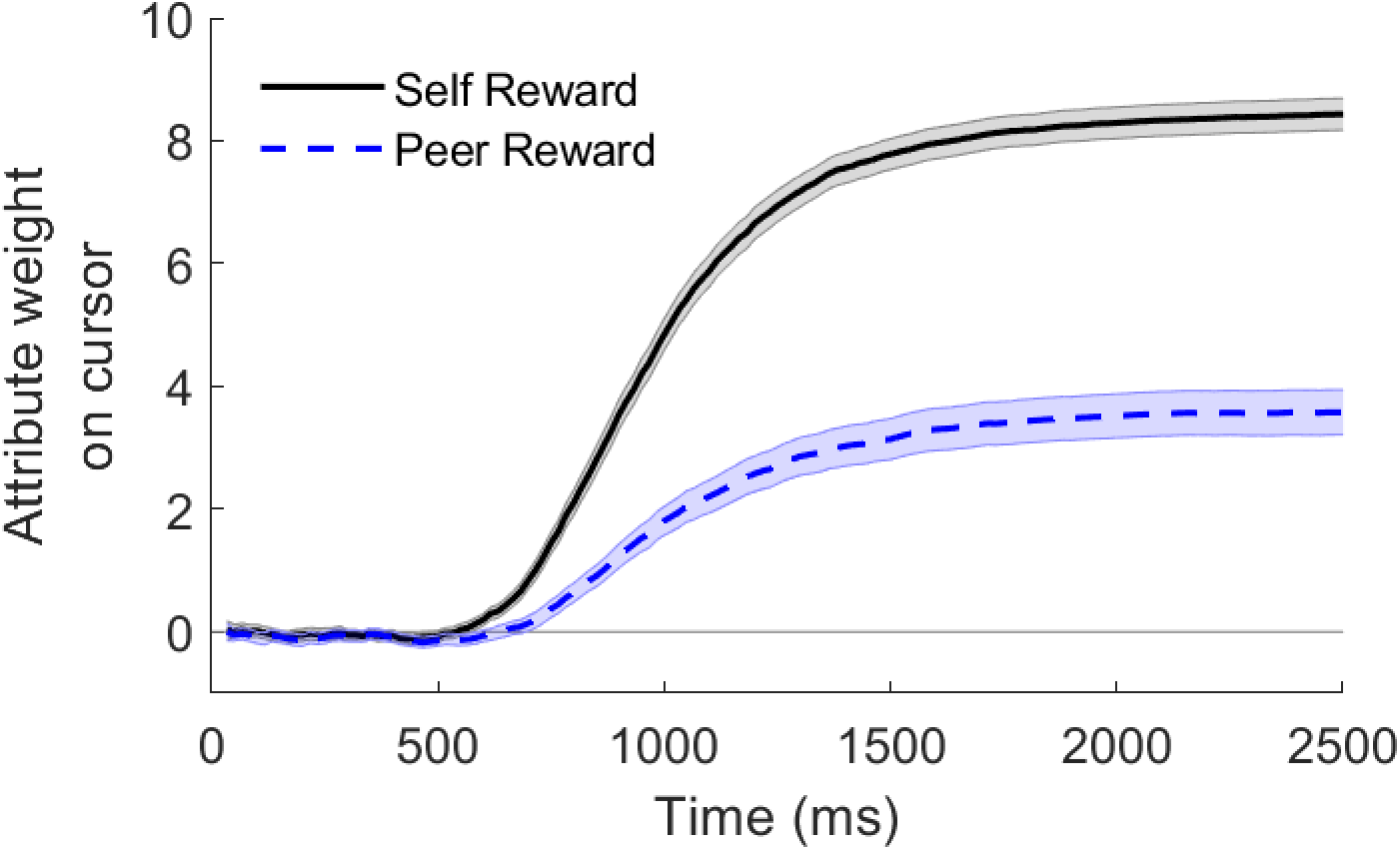
Influence of self and peer rewards on mouse trajectories. Results from the analyses exploring the mean effect of the relative self-reward of the right item over the left item (self_right_ – self_left_) and the relative peer rewards of the right item over the left item (peer_right_ – peer_left_), collapsed across all adolescent alone and watched conditions. Error bands denote standard errors.

**Fig. S3.**
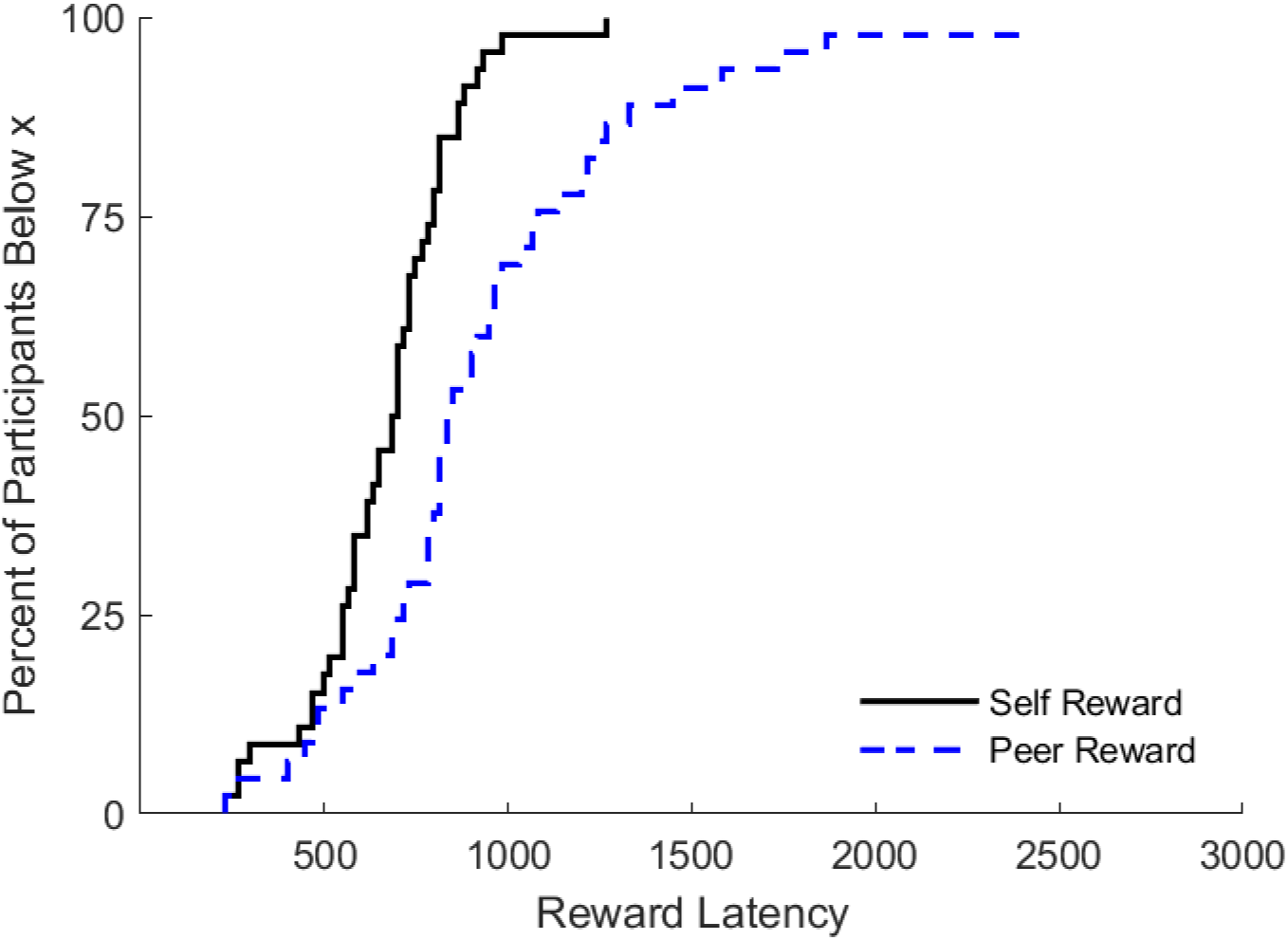
Results of reward latency estimation in the alone condition. The distribution of self and peer reward latencies are shown in this cumulative distribution graph for the watched condition.

**Fig. S4.**
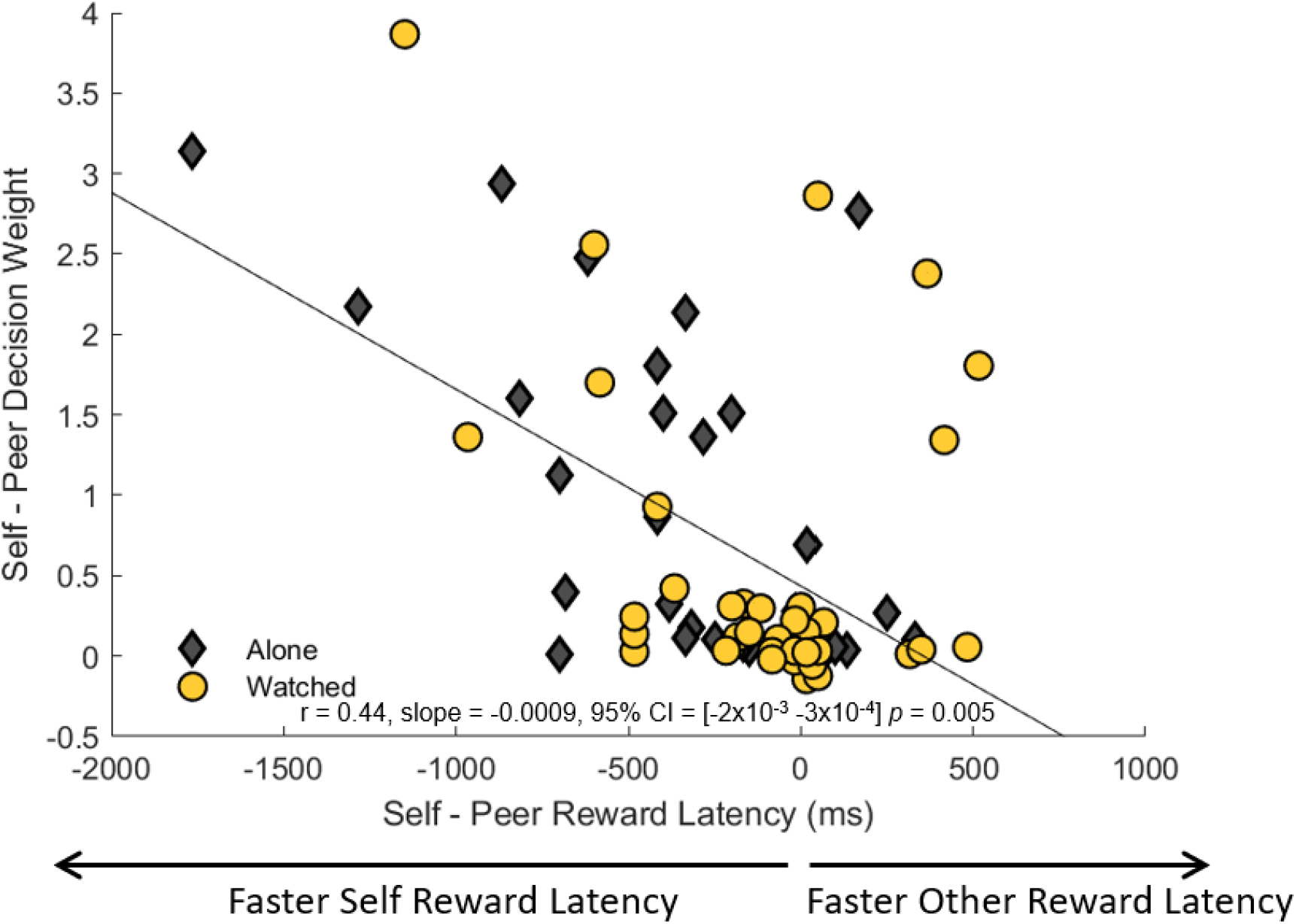
The relationship between the computational advantage of self-reward information and the relative weight adolescents placed on self- and peer-rewards. The markers depicts self, relative to peer, reward processing speeds as a function of this weighting difference, with grey diamonds representing the alone condition and yellow circles representing the watched condition. The line represents the best-fitting linear regression line.

**Fig. S5.**
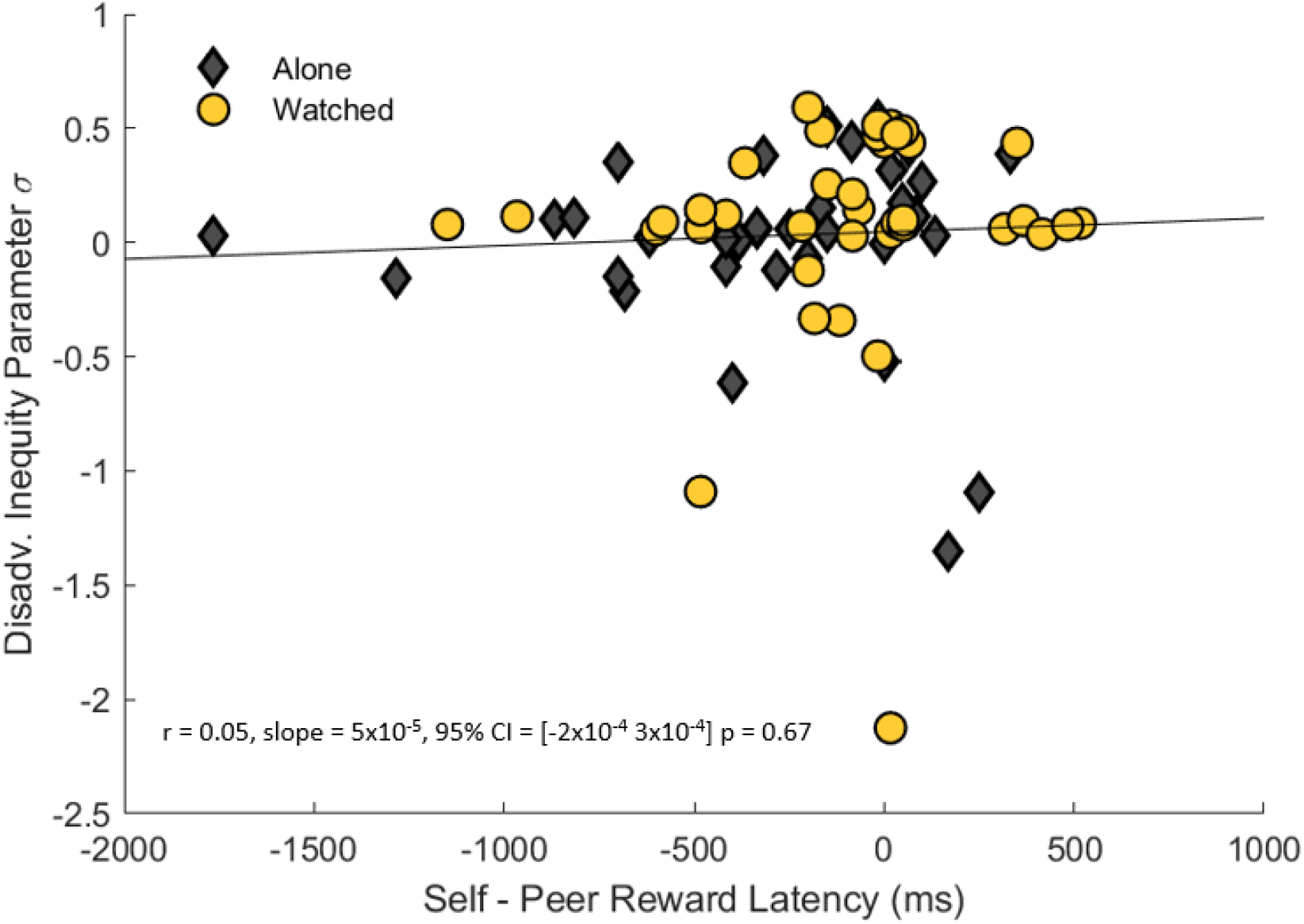
The relationship between disadvantageous inequity parameter σ and the computational advantage of self-reward information. The markers depict self, relative to peer, processing speeds as a function of individual best-fitting advantageous inequity parameter σ, with grey diamonds representing the alone condition and yellow circles representing the watched condition. The line represents the best-fitting linear regression line.

**Table S1.** Model AICs and BICs. Mean AIC and BIC for each of the four utility models in each participant type. Because the Charness-Rabin Model (Charness and Rabin 2002) had significantly lower AICs and BICs compared to the other utility models (paired t-tests, all *p*s < 0.001), we reported results of the Charness-Rabin Model to compare behavior between age groups and adolescent Alone and Watched conditions.

| **Model** | **Adults Alone** |  | **Adolescent Alone** |  | **Adolescents Watched** |  |
| --- | --- | --- | --- | --- | --- | --- |
|  | **AIC** | **BIC** | **AIC** | **BIC** | **AIC** | **BIC** |
| Fehr-Schmidt | 290.2 | 297.6 | 223.5 | 230.9 | 219.1 | 226.4 |
| Unbounded Fehr-Schmidt | 240.3 | 247.7 | 185.3 | 192.7 | 164.5 | 171.8 |
| Unbounded Fehr-Schmidt with peer parameter | 143.5 | 154.6 | 154.0 | 165.1 | 122.3 | 133.3 |
| Charness-Rabin | 140.4 | 147.7 | 149.6 | 157.0 | 118.2 | 125.5 |

